# Biomechanical Costs Influence Decisions Made During Ongoing Actions

**DOI:** 10.1101/2024.02.26.582113

**Authors:** Cesar Augusto Canaveral, William Lata, Andrea M Green, Paul Cisek

## Abstract

Accurate interaction with the environment relies on the integration of external information about the spatial layout of potential actions and knowledge of their costs and benefits. Previous studies have shown that when given a choice between voluntary reaching movements, humans tend to prefer actions with lower biomechanical costs. However, these studies primarily focused on decisions made before the onset of movement (“decide-then-act” scenarios), and it is not known to what extent their conclusions generalize to many real-life situations, in which decisions occur during ongoing actions (“decide-while-acting”). For example, one recent study found that biomechanical costs did not influence decisions to switch from a continuous manual tracking movement to a point-to-point movement, suggesting that biomechanical costs may be disregarded in decide-while-acting scenarios. To better understand this surprising result, we designed an experiment in which participants were faced with the decision between continuing to track a target moving along a straight path or changing paths to track a new target that gradually moved along a direction that deviated from the initial one. We manipulated tracking direction, angular deviation rate, and side of deviation, allowing us to compare scenarios where biomechanical costs favored either continuing or changing the path. Crucially, here the choice was always between two continuous tracking actions. Our results show that in this situation, decisions clearly took biomechanical costs into account. Thus, we conclude that biomechanics are not disregarded during decide-while-acting scenarios, but rather, that cost comparisons can only be made between similar types of actions.

**NEW & NOTEWORTHY:** In this study, we aim to shed light on how biomechanical factors influence decisions made during ongoing actions. Previous work suggested that decisions made during actions disregard biomechanical costs, in contrast to decisions made prior to movement. Our results challenge that proposal and suggest instead that the effect of biomechanical factors is dependent on the types of actions being compared (e.g., continuous tracking vs. point-to-point reaching). These findings contribute to our understanding of the dynamic interplay between biomechanical considerations and action choices during ongoing interactions with the environment.

## INTRODUCTION

Accurate interaction with our environment depends on our ability to continuously make use of a variety of sources of information. This includes not only external information about the spatial layout of our environment but also an implicit knowledge of our body’s biomechanical properties. For example, many studies of voluntary reaching have shown that when given a choice, humans tend to prefer actions that incur lower biomechanical costs over more effortful ones (1-6). For instance, decisions about which arm to use can be selectively biased by reducing the perceived effort associated with use of a given limb (7) as well as by the costs associated with additional forces applied on the limb by body motion (8). Biomechanical costs have also been shown to influence the selection of reach goals (1, 5, 9), in as little as 200ms after targets are presented (10). In fact, such costs have even been shown to bias perceptual decisions. In particular, when perceptual decisions were reported with reach actions, participants tended to reach to less effortful targets even if this led to decision errors (2, 4). One possible explanation is that during the process of action selection, a competition between potential movements unfolds within the sensorimotor system and is continuously biased by any relevant variables, including payoffs, costs, risks, etc. (11-14).

However, the studies mentioned above all involved situations in which decisions were made prior to the onset of movement. While this is common in laboratory studies, many real-life decisions are made while animals are already engaged in actions (walking through a crowd, playing a sport, or escaping predators in the jungle). Given that the nervous system evolved first and foremost to govern real-time interaction with the environment, it is important to examine to what degree the insights from “decide-then-act” paradigms translate to “decide-while-acting” situations (15-18). Indeed, some models argue that sensory information about the costs and benefits of an action and its dynamics influences the decision process even after movement begins (14), making it possible to adjust an initial action during ongoing movement, as a consequence of changing environments, sudden perturbations or motor costs (14, 15, 17, 19-24). Hence, the nervous system may track action-related variables such as motor cost and weigh decision-making in real time (25), a proposal that has been supported by various experimental studies (22, 24, 26, 27).

In a recent study, we investigated a range of factors that might influence arm movement choices in decide-while-acting scenarios (15). In the main experiment, participants tracked a continuously moving target with their hand and were sometimes presented with another target to which they could freely choose to switch. On different trials the alternative target was presented at different distances from the hand, at different angles with respect to the current tracking direction and had different sizes relative to the tracked target. As expected, all of these variables significantly influenced the decision to switch. What was unexpected, however, is that differences in the biomechanical costs associated with switching to a given new target versus another during continuous tracking did not appear to have any effect on switch decisions. More specifically, switching to a new target involved a point-to-point movement that always required more muscle torque than continuing to track a current target, but the cost in terms of torque difference between continuing to track the initial target versus switching to a new one varied significantly as a function of the tracking direction. Thus, if biomechanical costs are among the factors that influence choices, one would predict more decisions to switch in situations where the cost difference was smaller and fewer when it was larger. However, we saw no evidence for any dependence of switch choices on the tracking direction. This suggested that choice behavior during continuous tracking was remarkably insensitive to biomechanical costs. However, consistent with previous evidence from many “decide-then-act” experiments (1, 2, 4, 5), the biomechanical bias reemerged in a discontinuous version of the task in which participants tracked a target using point-to-point instead of continuous movements, such that when an alternative target was presented they always made a choice between two stationary targets. What could be the reason for this striking difference in the extent to which biomechanics influence choice behavior when executing different tasks?

One possible explanation for the absence of biomechanical biases during continuous tracking could be that the costs of alternative actions are simply ignored during decide-while-acting scenarios, perhaps because the systems sensitive to such costs are “busy” with the task of control. Another possibility could be that the constraints of tracking (staying in the target, matching velocity, etc.) outweighed the relatively minor and subtle differences in biomechanical costs. A third explanation might be that the two actions between which participants decided (i.e., continuous tracking versus point-to-point reaching to a new target) were sufficiently different that they are controlled by different circuits, so their relative costs could not be easily compared by the brain. This would also explain why the biomechanical bias was seen in the discontinuous task (in which the choice was between two point-to-point movements) and not in the continuous one.

Here, we test this third possibility using a decide-while-acting experiment in which the choice is always between two comparable continuous tracking actions. In the task, participants had to decide between continuing to track an initial target that moved along a straight path, versus tracking an alternative target that deviated from the initial direction along a curved trajectory. We selectively manipulated the initial tracking direction as well as the path deviation rate and direction (clockwise or counterclockwise) of the alternative target from trial to trial. This made it possible to compare between choice scenarios in which the biomechanical costs of continuing to track along a straight path were lower than those of following a new curved path and vice-versa. Some of these results have been previously reported in abstract form (28).

## METHODS

### Subjects and apparatus

Twenty right-handed subjects (11 women, 9 men) participated in the study. They all reported no known neurological or musculoskeletal disorders and had normal or corrected-to-normal vision. They all were naive to the purpose of the study and provided written informed consent before the experimental session was initiated. Each participant received a compensation of $15 CAD per hour for their participation. The protocol was approved by the Human Research Ethics Committee of the Faculté de Médicine, Université de Montréal.

Subjects sat in front of a 91-by-61 cm digitizing tablet (GTCO Calcomp IV, Columbia, MD) oriented in the horizontal plane, with a half-silvered mirror suspended 16 cm above and parallel to the digitizer. Visual stimuli were projected onto the mirror by an LCD monitor suspended 16 cm above, producing the illusion that the targets lie on the plane of the digitizing tablet. Each subject used their right arm to make movements while holding an upright digitizing stylus with their forearm in the semi-pronated position. The right shoulder was aligned to the center of the screen and the arm rested in a sling which supported the forearm just below the elbow. The sling length was adjusted so that the anchor point was approximately aligned with the subjects’ elbow when they held the stylus in the center of the screen (Figure 1A). The location of the stylus was sampled at 125 Hz with a spatial resolution of 0.013 cm. Kinematic data were stored and analyzed off-line.

**Figure 1.**
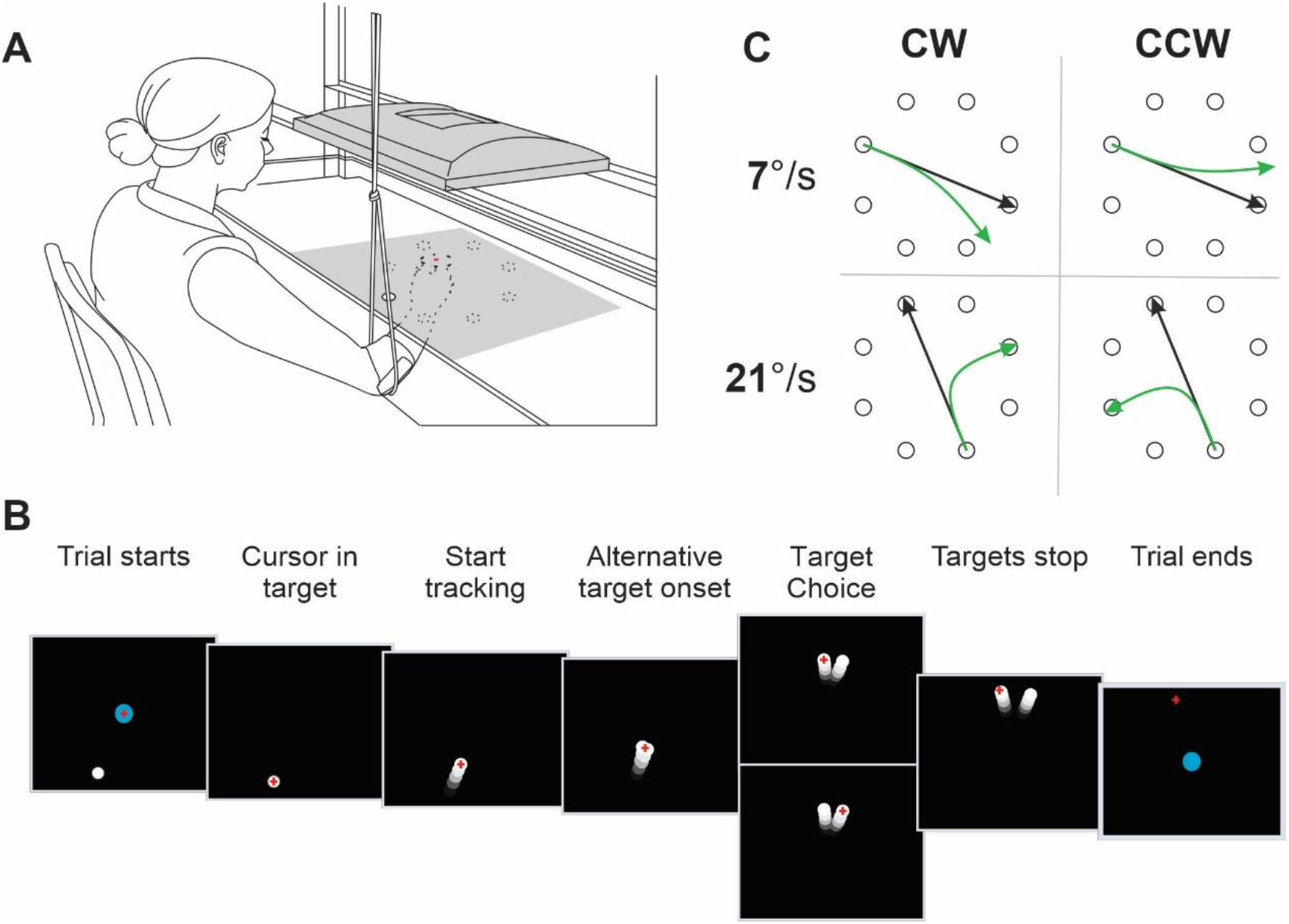
Experimental setup and task. A. Schematic of the experimental setup. The participant sat in front of a virtual environment where they had to perform a right-hand tracking task with their forearm supported with a sling to reduce friction. B. Task sequence. The trial started when the participant moved the cursor to the central starting position (blue circle, 2 cm), after which one of eight targets (white circle, 1 cm) appeared in the periphery indicating that the subject should reach to the target to start the tracking task. Once reached, the initial target started moving across the workspace. In some trials, as the target approached the center of the workspace, it split into two targets, one of which veered away from the initial straight-line trajectory. Participants were free to choose whether to continue tracking the initial target along its straight path (bottom “target choice” panel) or to change and track an alternative target along a curved path (top “target choice” panel). The trial ended when the targets reached the edge of the workspace, at which point they stopped moving and the next trial started. C. Layout of the targets in the workspace and combination of choice conditions. The alternative target could veer off the initial path clockwise (CW) or counterclockwise (CCW) at one of two angular deviation rates (slow, 7°/s; or fast, 21°/s). Example traces for tracking the initial target (black trace) vs. the alternative deviating target (green trace) are presented for reference.

### Protocol

The task sequence is depicted in Figure 1B. Subjects were instructed to track the position of a of moving target using a cursor (red crosshair; 1 cm line length) that represented the location of the stylus. Before the start of each trial, subjects were asked to bring the cursor to the center of the workspace. Subsequently, a target (white circle, 1 cm radius) appeared at one of eight positions equally distributed around an imaginary circle (12.5 cm radius, see Figure 1C for layout). The subject had to bring the cursor into the target, after which it began to move toward the center of the workspace at constant speed of 4.7 cm/s that was maintained as long as the cursor stayed within 1.5 cm of the target center. Otherwise, the target slowed down and stopped. This “initial target” moved in a straight path for 5 seconds. In most of the trials (80%), as the target approached the center of the workspace, an “alternative target” emerged from the initial target and diverged from it along a curved trajectory either to the right (i.e., clockwise) or to the left (i.e., counterclockwise) at one of two angular deviation rates (slow, 7°/s; or fast, 21°/s) for 4.3 seconds. The direction and speed of angular deviation were chosen pseudo-randomly. The subject was free to choose to either ignore the alternative target and continue tracking the initial target in the same direction (i.e., “straight path” trial, black traces in Figure 1B,C), or to track the alternative target and pursue the new path (i.e., “path change” trial, green traces in Figure 1B,C). Importantly, the tangential speed of the alternative target was always the same as that of the initial target (4.7cm/s) and therefore there was no advantage of choosing one over the other. At the end of the trial, subjects were required once again to bring the cursor to the center of the workspace for the start of the next trial. There were eight different tracking directions, that is 22°, 67°, 112°, 157°, 202°, 247°, 292°, and 337° (where 0° would be a movement to the right, orthogonal to the body’s midline). Subjects performed 40 practice trials before the experiment started and were allowed to take breaks between trials during the experiment if they wanted to. The experiment comprised 600 trials, 480 of which consisted of 15 repetitions of each combination of initial tracking direction, deviation direction and deviation rate (8 x 2 x 2, respectively) and an additional 120 trials which included 15 trials in each initial tracking direction for which no alternative target was presented. The experiment lasted approximately 1.5hrs.

### Biomechanical modeling

For each trial, we used a biomechanical model to estimate the net torque produced by the muscles during a trial. The model was built using the Simscape Multibody package within the Simulink simulation environment in MATLAB (Mathworks Inc. Natick, MA). The upper arm and forearm + hand limb segments were modeled for each individual subject as two thin rods with uniform mass distribution whose lengths and weights were estimated based on the subject’s weight and height (average males: upper arm, 33.4 cm, 2.3 kg; forearm + hand, 45.6 cm, 1.7 kg; average females: upper arm, 31.0 cm, 1.7 kg; forearm + hand, 42.3 cm, 1.3 kg) (29). The two limb segments were joined at the elbow with 1 rotational degree of freedom, and the proximal upper arm segment was joined to a static body with 1 rotational degree of freedom. The model was constrained to a two-dimensional (2-D) horizontal plane. The recorded position of the stylus was interpolated to 100 Hz using a 2-D spline and filtered at 20 Hz with a low-pass Butterworth filter (9^th^ order) with zero delay. Velocity was computed using a five-point differentiation routine (30), and then both position and velocity were up-sampled to 1,000 Hz via a process of linear interpolation and low-pass filtering to 20 Hz. Using inverse kinematics equations for a planar arm model, we calculated the angular position of each joint across time and then used an inverse dynamic model to calculate the associated muscle torques produced at the shoulder and elbow joints. We then calculated the net torque for each trial as the sum of the absolute muscle torques produced at both joints, averaged from the moment the cursor was brought into the initial target until the end of the trial. Note that because most trials involved smooth movements at a nearly constant speed (Figure 2A), and thus nearly constant torque, the average torque calculation was generally insensitive to the duration of the trial. As described in RESULTS, we found an anisotropy in the biomechanical cost of tracking across the different initial tracking directions. To quantify the biomechanical advantage of changing paths versus continuing in the same path in any given choice scenario, we calculated a metric of *path change cost* by dividing the average net torque of path change trials by that of straight path trials for each tracking direction. This metric allowed us to define “hard” and “easy” path change directions for every combination of deviation direction and deviation rate, as depicted in the outer bounds of the polar plots in Figure 2C and 3A.

**Figure 2.**
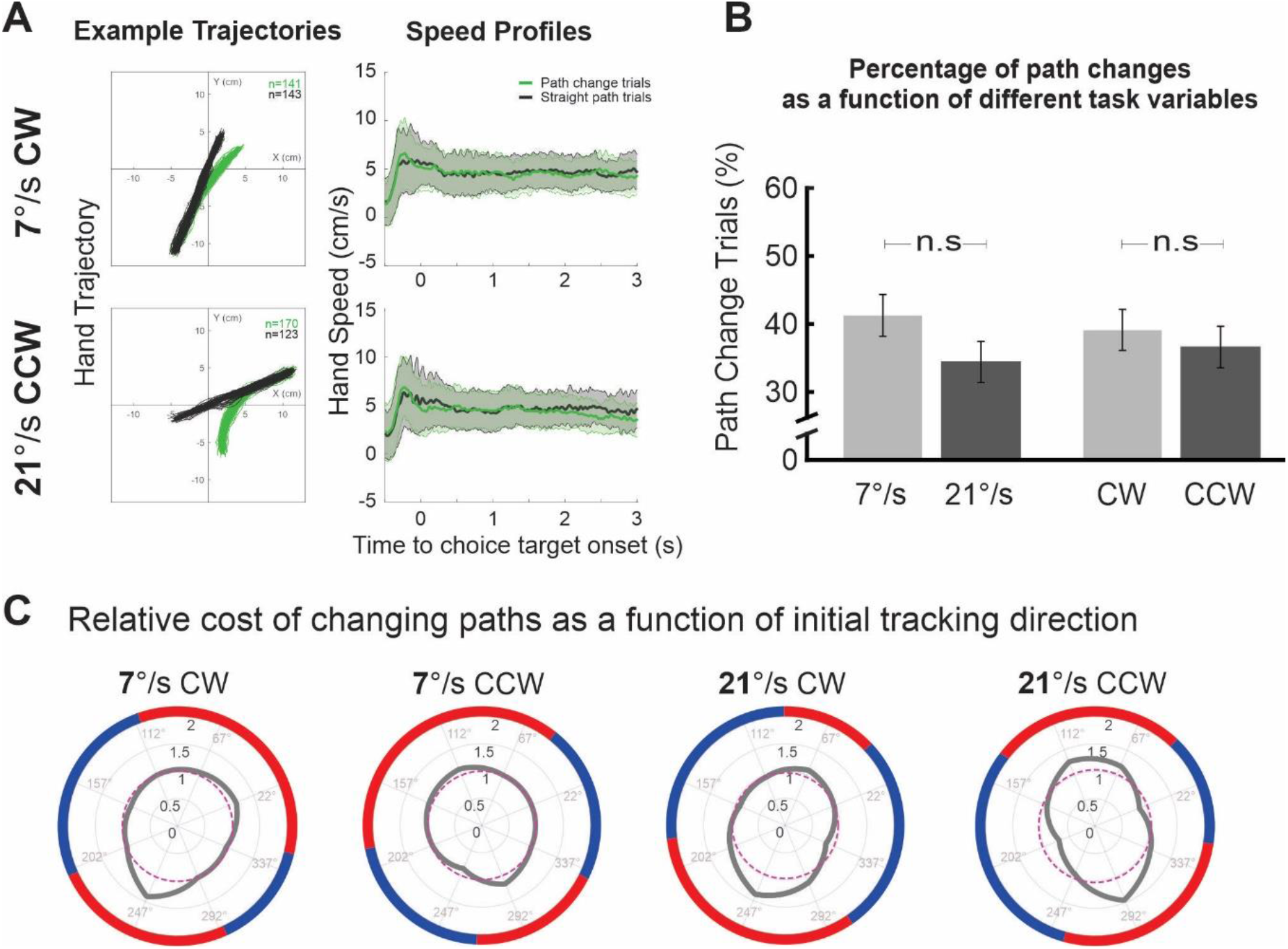
A. Trajectories and speed profiles for two example choice scenarios. As can be seen, participants completed the task accurately, with comparable speed profiles for path change (green trace) and straight path trials (black trace). Small colored numbers in the upper right trajectory plot represent the number of trials for that given trial outcome. B. Percentages of path change trials were not influenced by the rate nor the direction in which the alternative target veered off. Bar graphs and error bars represent medians and 95% confidence intervals, respectively. C. Relative cost of changing paths as a function of the initial tracking direction for all choice conditions. Polar plots represent the calculated torque ratio for changing paths versus continuing on the same straight path, as a function of the initial tracking direction (gray polygons) for each of the four deviation directions and rate combinations. The polar axis of the plot represents the eight different initial tracking directions used during the experiment. As expected, the relative cost of changing paths strongly depended on the initial tracking direction (significantly anisotropic, ꭕ2 above 51.74 and p<0.001 for all conditions). Purple dashed lines indicate the median relative torque for changing paths across all directions, allowing us to classify specific choice scenarios as “easy” (changing paths is less costly than not changing; blue border) or “hard” (changing paths is more costly than not changing; red border). CW, clockwise; CCW counterclockwise; n.s. not significant.

### Analysis of Choice Preference

We quantified the choice behavior as the percentage of path change trials for each initial tracking direction. In order to test whether the proportion of path change choices varied with the initial tracking direction, we performed a χ^2^ test over these percentages for every combination of deviation direction and rate. Kolmogorov-Smirnov tests revealed that the data were not normally distributed. We therefore reported data medians and used a non-parametric Wilcoxon rank sum test to compare choice behavior across the different conditions. Results were considered significant at *p* < 0.05.

## RESULTS

Twenty subjects performed a dual tracking task in which they had to choose whether to either continue tracking the initial target that moved along a straight path or to change paths to track an alternative target that deviated away from the initial tracking direction. Figure 2A shows examples of the cursor trajectory for a given tracking direction where “path change” and “straight path” trials (i.e., green and black traces, respectively) can be observed. The overall path change percentages for trials with slow deviation rates (i.e., 7°/s) versus fast deviation rates (i.e., 21°/s) were 41.17 % and 34.21 %, respectively. Similarly, on average, subjects changed their trajectories from the initial direction in 39.11 % of trials when the alternative target veered off to the right, and in 36.27 % of trials when it went to the left. Non-parametric 2 factor (deviation side; deviation rate) Friedman tests revealed no significant differences in the choice behavior between left and right deviations (χ^2^_(2, N = 32)_ = 0.35, *p =* 0.553), and a slight but non-significant trend toward more path changes for slower as compared to faster deviation rates (χ^2^_(2, N = 32)_ = 3.73, *p =* 0.053). Although there was no explicit reason for the subjects to change path, these results show that they did in fact do so quite often.

### Analysis of path change cost (net torque averages)

We determined the relative cost of changing path versus continuing along the initial one using a “*path change cost*” metric (see METHODS). When pooling all tracking directions together, there was no difference in relative torques between slow and fast deviation rates (median 1.07 vs. 1.06) and between right and left deviation directions (median 1.05 vs. 1.08) (all *p-values >* 0.05). Thus, we expect that any potential biomechanical effect that influenced choice behavior must be ascribed to the initial tracking direction. Indeed, as can be observed in Figure 2C, the calculated biomechanical cost of changing paths relative to continuing along the initial straight path strongly depended on the initial tracking direction. An analysis of uniformity revealed a significant anisotropy in the path change cost for every combination of deviation rate and direction (CW – slow: χ^2^_(7, N = 2324)_ = 53.65, *p* < .001; CW-fast: χ^2^_(7, N = 2333)_ = 52.49, *p* < .001; CCW – slow: χ^2^_(7, N = 2315)_ =51.74, *p* < .001; CCW-fast: χ^2^_(7, N = 2326)_ = 54.03, *p* < 0.001). This result allows us to classify specific choice scenarios as “easy” (changing paths is less costly than continuing straight) or “hard” (changing path is more costly than continuing straight) depending on the tracking directions for each combination (see METHODS).

### Analysis of choice preferences

Figure 3A shows the percentages of trials where there was a change of path as a function of the initial tracking direction for each choice target deviation rate and direction. The polar axes of Figure 3A panels represent the eight different initial tracking directions used during the experiment. As can be seen, subjects’ choices were different across directions. For example, in a scenario where the alternative target deviated CCW at a rate of 7°/s (Fig3A. second panel), subjects switched paths in 60.9% of trials when the initial tracking direction was heading to the left and slightly toward the subject (i.e., 200° ), whereas they changed paths in only 35.7% of trials when the initial direction was heading toward the subject and slightly to the right (i.e., 290°). This disparity in choice behavior across directions was quantitatively confirmed by an analysis of uniformity that revealed significantly non-isotropic patterns in the percentages of path changes across all initial tracking directions and for all combinations of deviation rate and direction (CW – slow: χ^2^_(7, N = 2324)_ = 67.85, *p* < 0.001 ; CW-fast: χ^2^_(7, N = 2333)_ = 74.27, *p* < 0.001; CCW – slow: χ^2^_(7, N = 2315)_ = 66.28, *p* < 0.001; CCW-fast: χ^2^_(7, N = 2326)_ = 80.54, *p* < 0.001). Importantly, the pattern of path change was congruent with biomechanical costs. As can be seen in Figure 3B, there is a higher proportion of path changes in choice scenarios with lower path change cost for most conditions (blue markers), as compared to those with higher path change cost (red markers) (CW – slow: 48.3 vs. 34.5 %, W_ranksum_ Z = 2.069, *p* = 0.038; CCW – slow: 40.0 vs. 33.3 %, W_ranksum_ Z = 1.753, *p* = 0.079; CW – fast: 41.4 vs. 20.7 %, W_ranksum_ Z = 3.214, *p* = 0.001; CCW – fast: 39.2 vs. 20.0 %, W_ranksum_ Z = 2.893, *p* = 0.004). This was further confirmed by a significant overall biomechanical bias when comparing percentages of path changes for “easy” vs. “hard” trials across all conditions (median 42.85 vs. 26.27 %; W_ranksum_ Z = 5.057, *p* < 0.001). In other words, subjects exhibited a significant preference for tracking targets along directions that incurred lower biomechanical costs.

**Figure 3.**
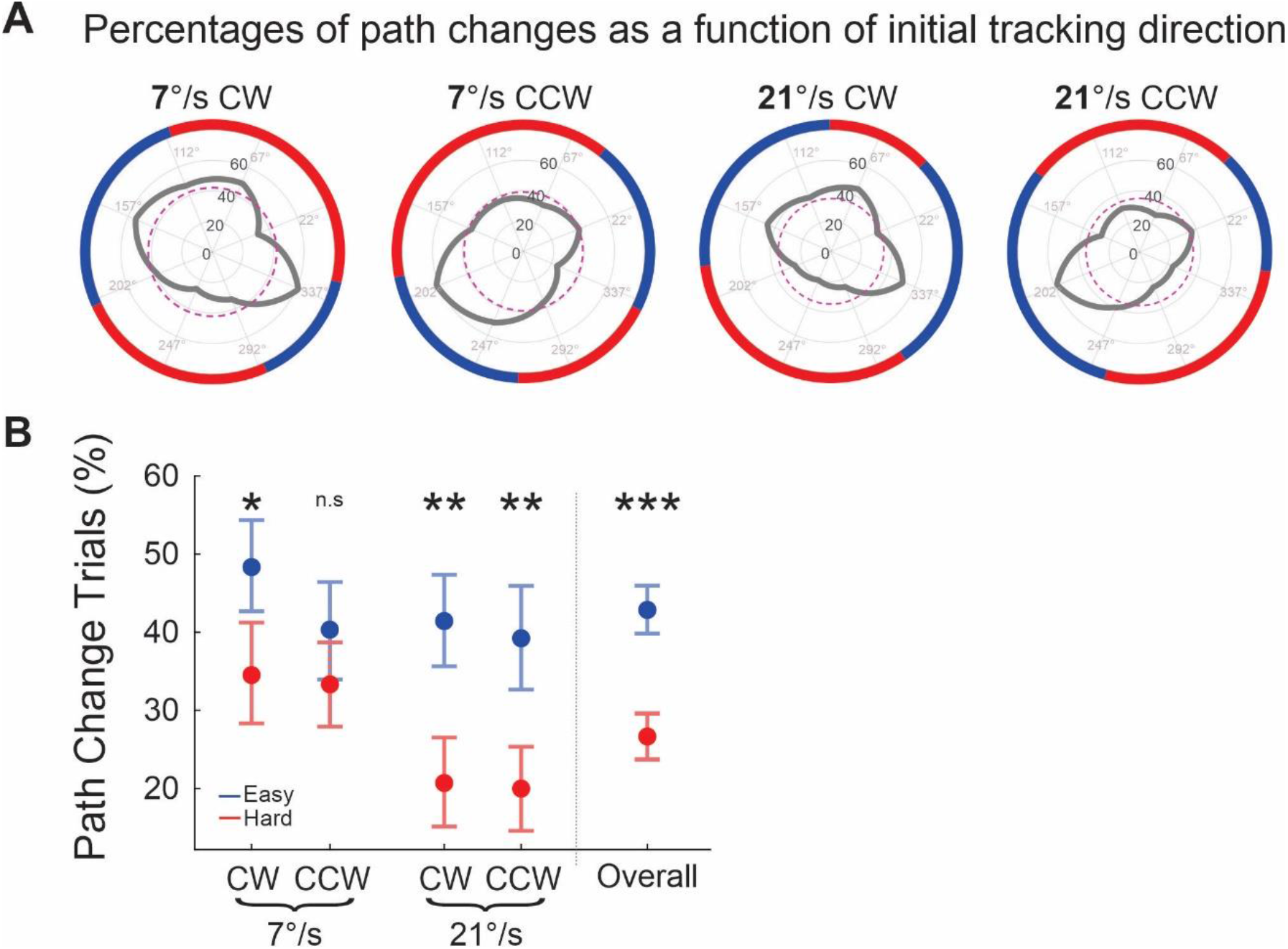
Path change percentages as a function of the initial tracking direction for all choice conditions. A. Polar plots represent the path change percentages as a function of the initial tracking direction (gray polygons). The polar axis of the plot represents the eight different initial tracking directions used during the experiment. As expected, the path change percentage strongly depended on the initial tracking direction (significantly anisotropic, ꭕ2 above 66.28 and p<0.001 for all conditions). Purple dashed lines indicate the median path change percentage across directions for every condition. Colored borders are imported from the torque ratio polar plots (Figure 2C), to facilitate the comparison of path change percentages with respect to the relative biomechanical cost in a specific condition. B. Comparison of easy versus hard path change trials revealed a higher proportion of path changes in choice scenarios where doing so incurred lower biomechanical cost compared to not changing paths and continuing in a straight line. This was true for most conditions. Low-cost trials are labeled as easy (blue markers) and high-cost trials are labeled as hard (red markers). Scatter plots and error bars represent medians and 95% confidence intervals, respectively. * p <0.05, ** p<0.01, and *** p<0.001; n.s. not significant; CW, clockwise; CCW counterclockwise.

### Changes of choice behavior over time

There is a possibility that the observed biomechanical bias could have been a consequence of the participants getting tired as the experiment progresses. To address this, we compared individual path change percentages between the first and last 80 trials of the experiment using a Friedman test that included the biomechanical effect as a factor to discriminate between low- and high-cost scenarios. After removing the effect of biomechanical bias, the analysis revealed a significant temporal effect between the early and late portion of the experiment (χ^2^_(2, N = 20)_ = 4.35, *p* = 0.037), suggesting a change in the participants’ choice behavior over the course of the experiment. Post hoc analysis revealed that this effect was selectively due to an increase in path changes in the low-cost scenarios (*p* < 0.001), with no difference across time in high-cost scenarios (p = 0.23; see Figure 4A). In other words, over the course of the experiment subjects might have specifically learned in which choice scenarios a path change resulted in lower biomechanical cost as compared to continuing to track the initial target.

**Figure 4.**
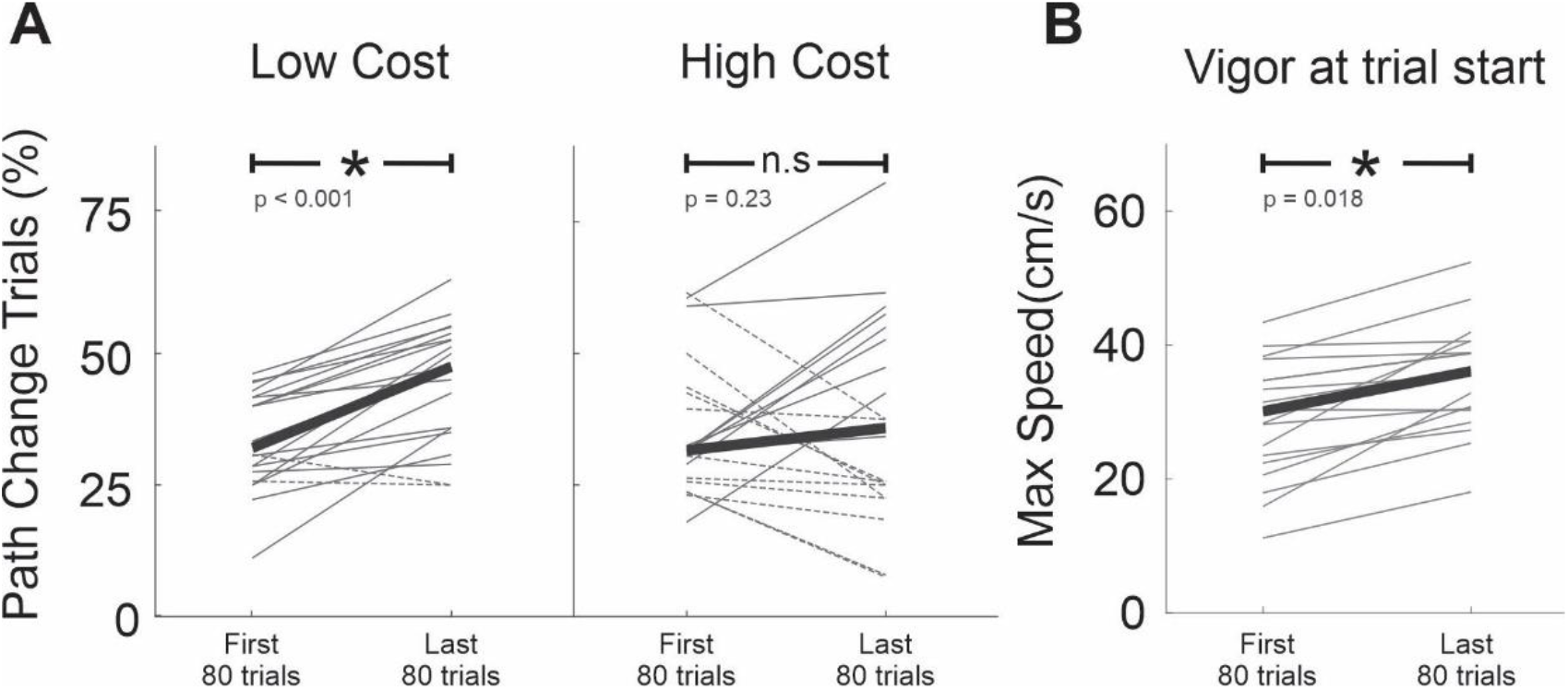
Path change percentage and participants’ vigor over time during the experiment. A. Comparison of path change percentages for low-cost and high-cost choice scenarios in the first and last 80 trials of the experiment. Individual subjects’ data are presented as thin gray lines. Thick black lines represent the median across subjects. Dashed lines indicate subjects exhibiting a reduction in path change percentages over time. B. Maximum speed from the center of the workspace into the initial target, used as a proxy for the initial movement vigor, during the first and last 80 trials of the experiment. n.s. not significant.

To examine whether this result could be explained by cumulative fatigue during the course of the experiment, we investigated the participants’ implicit motivation in performing the experiment. To do so, we calculated the maximum speed at which participants went from the central zone (blue circle in first panel of Figure 1B) into the initial tracking target and used this metric as a proxy for the participants’ vigor at trial start (31, 32). We then compared the overall maximum speed between the first and last 80 trials of the experiment using the Wilcoxon rank sum test. This analysis can be seen in Figure 4B. Surprisingly, results show that participants were significantly faster at starting the trial in the last 80 trials of the experiment compared to the first 80 trials (34.8 vs. 29.2 cm/s, respectively; W_ranksum_ Z = 2.367, *p* = 0.018). These results suggest that the changes in choice behavior might not be due to fatigue or decreased motivation, but rather to the participants’ knowledge about the biomechanical requirements of the task acquired through the course of the experiment.

## DISCUSSION

Humans and animals often find themselves in situations where they must make decisions between potential actions in the world around them – so called “embodied decisions”. In such circumstances, the potential actions are directly specified by sensory information about the geometry of the world and selection between them can take place through a biased competition between internal representations of potential movements (12, 13, 33). From this perspective, the selection between potential actions and the planning and control of these actions are functionally interdependent processes that influence each other in ways that take into account the properties of the actions and their required effort (1, 5, 34, 35).

Importantly, in many natural situations a decision must be made while one is already engaged in performing a particular action. In such a situation, one would expect the brain to compare the properties of the current action (including its biomechanical cost) with those of an alternative. Surprisingly, however, a previous study of deciding-while-acting found no consistent biomechanical influence on participants’ decisions between either continuing to track the same moving target or making a point-to-point movement to reach to a new one (15). This finding was at odds with evidence that subjects can take biomechanical costs into account when selecting between different point-to-point reaching movements (1-5, 9, 36).

To better understand this surprising result, here we tested the hypothesis that the actions between which participants decide must be sufficiently similar in order for the brain to be able to compare the effort required for each one and use that information to influence action decisions. We used a similar experimental strategy to the continuous task of Michalski et al. (15), except that both potential actions (i.e., tracking the initial target or the alternative one) involved continuous manual tracking. Thus, the movement choices were comparable in terms of their kinematic and dynamic properties as well as in terms of the control circuits that might govern their execution. Although there was no explicit reason why participants would choose the curved path over the straight one, results revealed that they did so quite often, allowing us to quantify the motor cost of either choice and to determine its influence on the choice behavior.

Our main finding is that when choosing between two continuous tracking movements, participants chose to follow and track targets whose path headed in directions that incurred lower biomechanical costs, even if that implied veering off and changing from a predictable straight initial path. This suggests that the brain is capable of computing biomechanical costs even during ongoing movement execution, and that these costs are significant enough to bias choice behavior. Importantly, this effect appears not to be caused by participants getting fatigued, but instead becomes stronger over time, perhaps because participants learned which options would incur the lowest effort as the experiment progressed.

In our experiment, both alternative actions involved simultaneously matching the direction and velocity of the participants’ hand to that of the tracked targets (37-39). Since the targets moved at similar speeds (see *speed profiles in* Figure 2A), and since there were no explicit incentives favoring either option, participants likely based their decision on minimizing the biomechanical cost associated with the action (40). This differs significantly from the type of decision that participants encountered in the “*continuous task”* of our previous study (15). In that scenario, participants had to decide between continuing to track versus making a point-to-point movement to a new target – two movements with very different control policies, which may have prevented a comparison between their respective energy expenditure. In contrast, in a “*discontinuous task”* in which decisions were made between two point-to-point movements, participants did favor low-effort targets, highlighting the increased role of biomechanical factors in decision-making when the task requirements were comparable.

An alternative explanation for the absence of biomechanical cost effects in our previous study (15) is that the magnitude of the cost difference between tracking and one point-to-point movement versus tracking and another point-to-point movement was not large enough. Perhaps the biomechanical influence on decisions might have been higher if the cost differences between options, in proportion to the cost of the ongoing action, were larger, consistent with Weber’s law. However, this interpretation is unlikely because the cost of choosing to make a given point-to-point movement in that study varied considerably from being 2 to 3 times larger than the cost of continuing to track (Figure 3A of Michalski et al. (15)) but this had no impact on choice. In contrast, in our study cost differences associated with choosing to switch targets vs continuing along the same path varied by less than 12% (Figure 2C), and yet biomechanical cost effects were observed.

Our results are compatible with previous studies suggesting that whole-body dynamics influence decisions made during actions. For example, a study conducted by Grießbach et al. (2021) (17) reported that decisions about which side to walk around an obstacle are largely influenced by the anticipated effort associated with each option given their ongoing motion. Indeed, participants preferred avoiding an obstacle by doing a lateral step (i.e., less effortful option) rather than a cross-over step, sometimes even at the expense of gaining a higher reward (17). Similar biases have also been reported in a computer mouse tracking task (41), where participants most often chose movements with lower amplitude even if that option sometimes resulted in lower rewards (41, 42). It would be interesting to know whether their results would hold if the dynamics of alternative movements had been different.

Taken together, these results are consistent with the notion of a functional co-dependance between the neural mechanisms of action selection and their specification, which makes sense given that the nervous system evolved to govern our interactions with the environment (43-45). Many studies have suggested that decisions between actions involve the specific frontoparietal circuits implicated in controlling those actions – including dorsal premotor and medial parietal cortex for reaching (46-49), ventral premotor and anterior intraparietal cortex for grasping (50), and frontal eye fields and lateral intraparietal cortex for saccades (13, 51, 52). If decisions between a given type of action involve competition between cells within a given circuit specialized for that type of action, then that competition might be biased by the relative effort of the choices (as we observed in this study). But if the decision is between two different kinds of actions, then the effort costs may not factor into the choice (as observed by (15)) because they are represented in distinct circuits.

Although continuous hand-tracking is known to involve similar frontoparietal circuits as those involved in point-to-point hand movements, additional circuits are implicated in integrating visual information over time and coordinating motor adjustments. For example, manual tracking involves smooth pursuit eye movements while point-to-point reaching involves saccades, and the networks that control these different eye movements involve largely different circuits and/or non-overlapping cell populations (53-58). A recent study by Coudiere and Danion (59) highlighted the different movement dynamics associated with continuous tracking of a moving target compared to making a point-to-point reach to a static one. By progressively increasing the frequency with which a moving target appeared during a visuomotor tracking task, the authors found that participants transitioned from performing point-to-point reaches to continuous hand tracking quite abruptly when the frequency of target appearance exceeded 3Hz (59). Their results, along with similar conclusions made by others (60, 61), suggest that continuous manual tracking of a moving target and point-to-point reaches to a static one involve different sensory processing and different sources of information about errors (position plus velocity versus just position) (38). It is therefore possible that the recruitment of different subsets of regions for continuous versus point-to-point manual actions makes it difficult for the brain to fully compare their costs and benefits. Interactions between these systems and the factors that influence decisions within and between the associated circuits will be an important topic for future studies.

## ACKNOWLEDGEMENTS & FUNDING

The authors are supported by the Natural Sciences and Engineering Research Council of Canada (RGPIN-05345) and (RGPIN-05408), by the *Fonds de Recherche du Québec – Santé* (FRQS # 288624), and by the FRQNT Strategic Clusters Program (2020-RS4-265502 - Centre UNIQUE - Union Neurosciences & Artificial Intelligence - Quebec).

## CONTRIBUTIONS

CAC and WL collected and analysed the data, CAC prepared the figures. CAC, AG, and PC conceived the experiment and drafted the manuscript.

